# Insulin Signaling Attenuates GLUT4 Endocytosis in Muscle Cells *via* GSK3α-Dyn2-Bin1 Interplay

**DOI:** 10.1101/2021.08.23.457303

**Authors:** Jessica Laiman, Julie Loh, Wei-Chun Tang, Mei-Chun Chuang, Bi-Chang Chen, Yi-Cheng Chang, Lee-Ming Chuang, Ya-Wen Liu

## Abstract

Insulin-induced translocation of glucose transporter 4 (GLUT4) to the plasma membrane of skeletal muscle is critical for postprandial glucose uptake; however, whether the internalization of GLUT4 into cells is also regulated by insulin signaling remains unclear. Here, we discover that the activity of dynamin-2 (Dyn2), pivotal GTPase catalyzing GLUT4 internalization, is regulated by insulin signaling in muscle cells. The membrane fission activity of Dyn2 is inhibited in muscle cells through binding with the SH3 domain-containing protein Bin1. Phosphorylation of Serine848 on Dyn2 by GSK3α or the mutations of Bin1-SH3 in patients with centronuclear myopathy, elevate the activity of Dyn2 due to reduced binding affinity toward Bin1. The augmented Dyn2 fission activity in muscle cells leads to GLUT4 internalization and Bin1-tubule vesiculation. Together, our findings reveal a new role of insulin signaling in glucose metabolism and muscle physiology *via* attenuating Dyn2 activity thus regulating GLUT4 endocytosis in muscle cell.

## Introduction

Skeletal muscle is the primary responsive tissue for insulin-stimulated glucose uptake, and its insulin resistance often presages the development of type II diabetes (Kraegen et al., 1985; Lillioja et al., 1988; Warram et al., 1990). The insulin-induced glucose uptake is executed by the glucose transporter GLUT4, which is specifically expressed in muscle and adipose tissues (Birnbaum, 1989; Charron et al., 1989; James et al., 1989). Upon insulin stimulation, the insulin receptor-phosphoinositide 3-kinase (PI3K)-Akt signaling cascade is initiated to mediate translocation of GLUT4 from intracellular membrane compartments to the cell surface, thus reducing blood glucose (Beg et al., 2017; Katome et al., 2003; Leto and Saltiel, 2012). Upon insulin withdrawal, GLUT4 is internalized back into the cell through dynamin-dependent endocytosis (Antonescu et al., 2008; Kao et al., 1998; Volchuk et al., 1998). Contrary to its role in promoting GLUT4 exocytosis, the role of the insulin signaling pathway in GLUT4 endocytosis remains elusive.

The internalization of GLUT4 is mainly through dynamin-2 (Dyn2)-mediated endocytosis both in muscle and adipocytes. Dyn2 is a large GTPase catalyzing membrane fission at the last step of clathrin-mediated, caveolae-dependent, as well as clathrin and caveolae-independent (CCI) endocytosis (Ferguson and De Camilli, 2012; McMahon and Boucrot, 2011; Mettlen et al., 2009). Dynamin consists of five functional domains: the N-terminal GTPase domain binds and hydrolyzes GTP, the middle domain and GTPase effector domain (GED) together form the stalk region for self-assembly and oligomerization, the pleckstrin homology (PH) domain binds phosphatidylinositol-4,5-bisphosphate (PI(4,5)P_2_) at the plasma membrane, and the C-terminal proline-arginine rich domain (PRD) binds SH3 domain-containing proteins (Fig 1A) (Chappie et al., 2010; Faelber et al., 2011; Grabs et al., 1997; Zheng et al., 1996). In addition to recruiting Dyn2 to clathrin-coated pits (Daumke et al., 2014; Meinecke et al., 2013; Taylor et al., 2011), SH3 domain-containing interacting partners have also been shown to positively or negatively regulate the fission activity of dynamin (Boucrot et al., 2012; Farsad et al., 2001; Neumann and Schmid, 2013; Takei et al., 1999; Yoshida et al., 2004).

**Figure 1.**
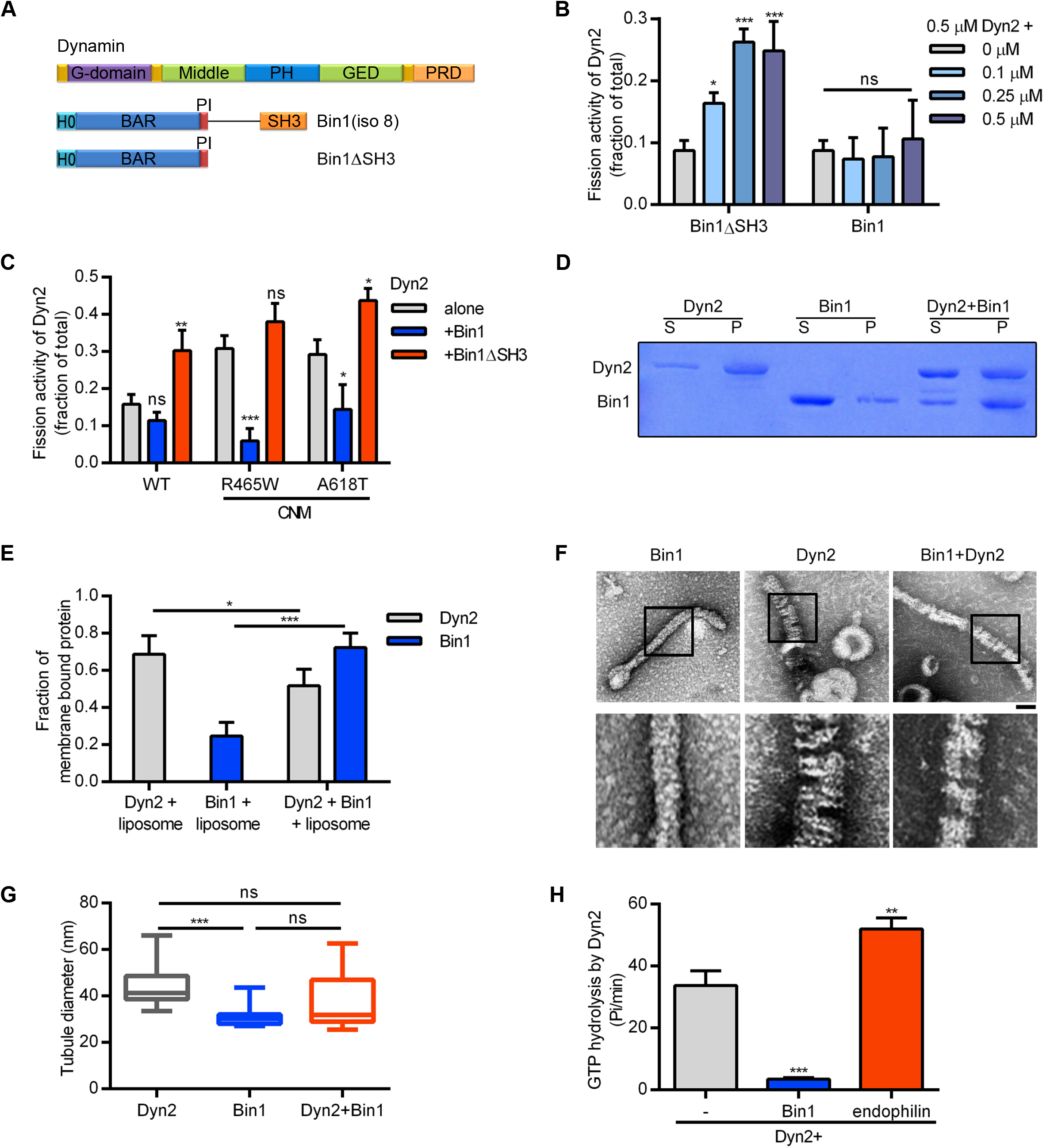
Bin1 inhibits Dyn2 fission activity through SH3 domain. (A) Domain structure of Dynamin and Bin1 constructs used in this study. (B) *In vitro* fission assay of Dyn2. 0.5 μM purified Dyn2 was incubated with GTP and SUPER template in the presence or absence of indicated Bin1 proteins. The fission activity was measured as the amount of vesicle released from the SUPER template (n = 3). (C) Effect of Bin1 on the fission activity of Dyn2 mutants (n = 3). (D) Liposome binding ability. 0.5 μM purified Dyn2 or Bin1 were incubated with 150 μM, 100 nm liposome for 10 min at 37°C. Liposome-bound proteins (pellet, p) were separated from unbound ones (supernatant, s) with centrifugation sedimentation. The ratio of liposome-bound proteins were quantified and shown in (E) (n = 3). (F) Assembly of Dyn2 or Bin1 on liposome. 1 μM Dyn2 and/or Bin1 were incubated with liposome and then visualized with negative stain TEM. Scale, 100 nm. Boxed areas were magnified and shown as below. The diameter of tubulated membrane were quantified and shown in (G) (n ≥ 11). (H) Liposome stimulated GTPase activity of Dyn2. 0.5 μM Dyn2 was incubated with 150 μM liposome with 0.5 μM Bin1 or Endophilin in the presence of 1 mM GTP at 37°C. The rate of GTP hydrolysis was measured using a colorimetric malachite green assay (n = 3). Bar graphs are shown as average ± SD, box plots are shown as median ± min and max value. Data are analyzed with one-way ANOVA. ns, no significance; *P< 0.05; **P< 0.01; ***P< 0.001.

Dyn2 is ubiquitously expressed in mammals, yet mutations of Dyn2 lead to tissue-specific diseases, one of which is autosomal dominant centronuclear myopathy (CNM) (Bitoun et al., 2005; Zhao et al., 2018). CNM is a congenital muscle disease characterized by centralized nuclei, myofiber atrophy and progressive muscle weakness mainly caused by mutations of three somatic genes, *MTM1*, *DNM2* and *BIN1* (Cowling et al., 2012). Previous studies have demonstrated that CNM-associated mutants of Dyn2 are hyperactive due to abnormal hyper-assembly (Chin et al., 2015; Kenniston and Lemmon, 2010; Wang et al., 2010). Expression of these Dyn2 mutants causes T-tubules, the specialized structures derived from sarcolemma invaginations, to be abnormally shaped in mice (Cowling et al., 2011) and zebrafish (Gibbs et al., 2014), and even become fragmented in *Drosophila* (Chin et al., 2015), suggesting defect in T-tubule maintenance as one of the pathological mechanisms of CNM.

Besides Dyn2, mutations of its binding partner, Bin1/amphiphysin 2, also result in CNM (Bohm et al., 2014; Nicot et al., 2007). Bin1 is an N-BAR and SH3 domain-containing protein whose isoforms are either ubiquitously or tissue-specifically expressed (Hong et al., 2014). N-BAR domains is a subset of the BIN-Amphiphysin-Rvs (BAR) domain superfamily that encodes an N-terminal amphiphatic helix (H0, Fig 1A). N-BAR forms a banana-shaped homodimer that binds membrane for curvature sensing and generation, while the SH3 domain binds the PRD of dynamin (Owen et al., 1998; Peter et al., 2004). Intriguingly, the muscle isoform of Bin1 is equipped with a unique region, PI motif, made up of 15 highly basic amino acid residues important for PI(4,5)P_2_ binding (Lee et al., 2002). Bin1 localizes to T-tubules and is required for their biogenesis in skeletal muscle (Al-Qusairi and Laporte, 2011; Lee et al., 2002; Razzaq et al., 2001). A splicing-defective Bin1 mutation lacking the PI motif also results in CNM (Böhm et al., 2013). Considering Dyn2 and Bin1 are interacting partners with opposing effects on T-tubule, severing and generating it respectively; it is tempting to hypothesize that a common molecular pathway regulated by these two proteins underlies CNM pathology.

Several animal model studies have shown that the downregulation of Dyn2 could serve as a therapeutic approach not only for Dyn2-related CNM (Buono et al., 2018; Rabai et al., 2019), but also to rescue phenotypes of Bin1- and MTM1-related CNM mouse models (Cowling et al., 2014; Cowling et al., 2017; Tasfaout et al., 2017). These findings suggest a role for Bin1 as negative regulator of Dyn2, yet the molecular and regulatory mechanism are unclear. Here, we uncover the mechanism of Dyn2 inhibition by Bin1, and find that this inhibition could be relieved by CNM-related mutants of Bin1 with truncated SH3 domain or by posttranslational modification through GSK3α phosphorylation on Dyn2-PRD. This regulation works to ensure T-tubule maintenance and control GLUT4 endocytosis, accentuating the importance of Dyn2 and Bin1 for proper physiological function and insulin-dependent GLUT4 trafficking in skeletal muscle.

## Results

### Dual functions of Bin1 in regulating Dyn2-mediated membrane fission

To investigate the effect of Bin1 on Dyn2 fission activity, we utilized an *in vitro* membrane fission assay to measure Dyn2-catalyzed vesicles released from supported bilayer with excess membrane reservoir (SUPER) templates (Neumann et al., 2013; Pucadyil and Schmid, 2010) (Fig 1B). As previously shown for the N-BAR domain of endophilin (Neumann and Schmid, 2013), we found that the N-BAR domain along with PI motif of Bin1 (Bin1ΔSH3) enhanced Dyn2-mediated membrane fission in a concentration-dependent manner. In contrast, full-length Bin1 had no effect (Fig 1B). To test whether Bin1 could inhibit the fission activity of dynamin, we checked its effects on membrane fission by Dyn1 (Fig S1A) and CNM mutants of Dyn2, which possess higher fission activity than WT-Dyn2, due to stronger curvature generation and hyper self-assembly, respectively (Chin et al., 2015; Liu et al., 2011). As expected, these proteins exhibited greater fission activity than Dyn2, which importantly, was inhibited in the presence of Bin1 (Fig S1A-C and 1C). These results show the dual roles of Bin1 on Dyn2 fission activity in which enhancement by its N-BAR domain and suppression through its SH3 domain, together produce a net negative regulatory effect on Dyn2 membrane fission activity.

To test if the inhibitory effect of Bin1 was due to alteration of membrane binding ability, we performed liposome binding assays. We observed a small, but significant reduction of Dyn2 binding in the presence of Bin1 and, interestingly a significant increase in Bin1 membrane binding (Fig 1D-E, and S1D). These data suggest that Dyn2 relieved Bin1 autoinhibition (Kojima et al., 2004; Wu et al., 2014) to improve Bin1 binding to membrane and that membrane-bound Bin1 inhibits the activity of membrane-bound Dyn2. To visualize how Dyn2 and Bin1 assemble on membranes, we next used negative-stain transmission electron microscope (TEM) to image Dyn2 and Bin1 proteoliposomes. Dyn2 assembly on 100 nm-liposomes resulted in tubule formation with average diameter of 44.5 ± 2.3 nm and a defined spiral-like pattern (pitch = 8.46 ± 2.31 nm) (Fig 1F-G and S1E). Assembly of Bin1 on liposomes also formed membrane tubules but with indiscernible patterning and smaller diameter (31.4 ± 1.2 nm). Co-incubation of Dyn2 and Bin1 led to fewer membrane tubules with less-ordered, spiral-like protein assembly (pitch = 18.35 ± 8.10 nm) and intermediate tubule diameter (37.4 ± 3.7 nm), indicating altered protein assembly on membrane (Fig 1G and S1E). Consistent with its inhibition on Dyn2-mediated membrane fission, Bin1 also suppressed Dyn2 GTPase activity in the presence of liposomes (Fig 1H and S1F). It is worth noting that another SH3 domain-containing Dyn2-binding protein, endophilin, promoted liposome-stimulated GTPase activity of Dyn2, indicating the different SH3 domain binding partners exert distinct regulatory effects toward Dyn2 (Fig 1H). Together, these results indicate that Bin1 negatively regulates Dyn2 fission activity via binding to and altering assembly of membrane-bound Dyn2.

### PI motif of Bin1 does not affect membrane fission activity of Dyn2

Muscle specific isoforms of Bin1 contain a unique positively-charged, 15 amino acid stretch called the PI motif, which enhances the lipid binding ability of Bin1 (Lee et al., 2002). Given that membrane-bound Bin1 inhibits Dyn2 activity, we tested the potential role of PI motif in Dyn2-mediated membrane fission assays using different Bin1 variants (Fig S2A). To our surprise, PI motif did not affect Bin1 regulation on Dyn2 fission activity (Fig S2B). By contrast, the binding affinity between Bin1 and Dyn2 was increased by the truncation of PI motif probably due to the reduction of autoinhibition (Fig S2C-D, (Kojima et al., 2004; Wu et al., 2014)). Considering that Bin1 is important for the formation of T-tubules, which are enriched with cholesterol and sphingomyelin resulting in higher membrane rigidity (Rosemblatt et al., 1981; Roux et al., 2005), we added cholesterol and different percentages of brain sphingomyelin (BSM) to our lipid template. Even on these stiffer membranes, the PI motif did not significantly affect Dyn2 fission activity (Fig S2E). Nonetheless, and consistent with a previous report (Lee et al., 2002), we found that the PI motif significantly improved the binding of Bin1 N-BAR domain to membrane (Fig S2F and G). In accordance with its ability to promote membrane binding, the PI motif gave rise to greater membrane tubulation of SUPER templates *in vitro* (Fig S2H and I) and of plasma membrane in cells (Fig S2J). These results suggest that the PI motif is dispensable for regulating Dyn2 fission activity but important for membrane binding and tubulation of Bin1.

### Enhanced Dyn2 fission activity by CNM-associated mutants of Bin1-SH3

Among CNM-related Bin1 mutations, two of them (Fig 2A, Bin1^Q434X^ and Bin1^K436X^) generate premature stop codons that cause truncation of Bin1 SH3 domain (Bohm et al., 2010; Nicot et al., 2007). We thus proposed the pathological mechanism of these CNM-associated Bin1 mutants might be due to the loss of SH3-PRD inhibition. Membrane binding and tubulation ability of both Bin1^Q434X^ and Bin1^K436X^ *in vitro* were comparable to that of wild type Bin1 (Fig S3A and 2B). Under TEM, assembly of Bin1^WT^, Bin1^Q434X^ and Bin1^K436X^ on liposome resulted in the formation of membrane tubules with indistinguishable patterning (Fig S3B). Using GST-tagged PRD domain of Dyn2 (GST-PRD) to pull down Bin1 protein, we observed a small but significant decrease in binding of Bin1^Q434X^ and Bin1^K436X^ to GST-PRD, compared to Bin1^WT^ (Fig 2C-D). We then investigated whether this change in SH3-PRD binding affinity would affect Bin1 regulation of Dyn2 fission activity. Remarkably, Bin1^Q434X^ and Bin1^K436X^ behaved in a similar way to Bin1ΔSH3 (Fig 1B) in that they elevated Dyn2 fission activity (Fig 2E). Intriguingly, the binding affinity between Dyn2 and Bin1^K436X^ is only slightly reduced from 180 ± 60 nM to 267 ± 70 nM (Fig 2F-G). This result suggests that weakening SH3-PRD interaction could serve as a mechanism to relieve Bin1-SH3 inhibition and allowing the stimulatory activity of the N-BAR domain to become the predominating effect of Bin1 on Dyn2 fission activity. Incubation of Bin1^Q434X^ or Bin1^K436X^ together with Dyn2 on liposomes led to the formation of more abundant membrane tubules with slightly smaller diameter on average than those derived from the co-assembly of wild type Bin1 and Dyn2 (Fig 2H and I).

**Figure 2.**
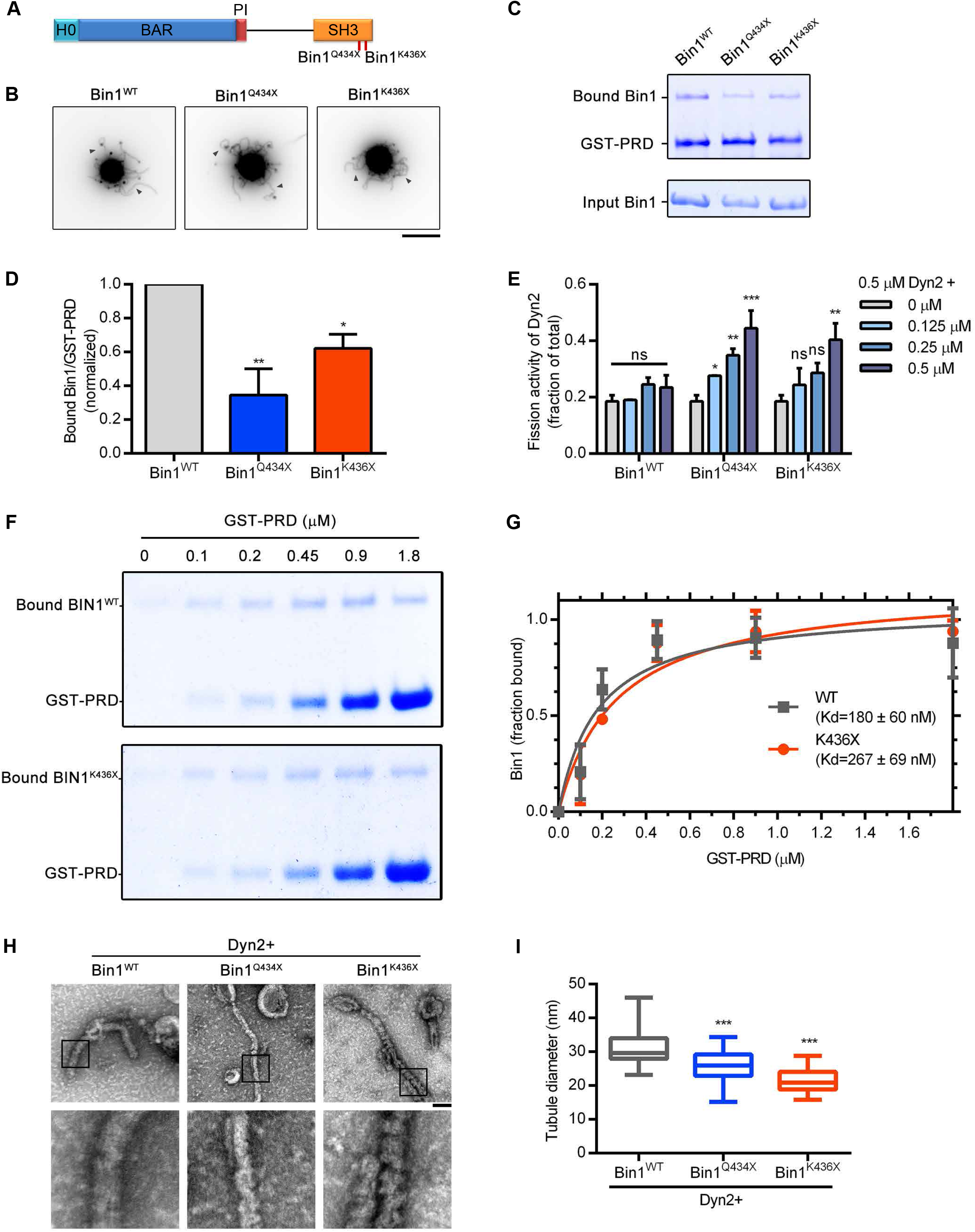
Effects of CNM-Bin1 on Dyn2 fission activity in vitro. (A) CNM-associated Bin1 mutants used in this study. (B) Membrane tubulation ability of Bin1 mutants. 0.5 μM wild-type and mutant Bin1 were incubated with SUPER templates for 10 min at room temperature and imaged under fluorescent microscopy. Black arrow heads indicate the tubulated membrane. Scale, 5 μm. (C) Binding ability between Bin1 and the PRD of Dyn2. GST or GST-PRD was incubated with indicated purified His-tagged Bin1, and then washed with PBS. PRD-bound Bin1 was analyzed with SDS-PAGE and Coomassie Blue staining. The ratio of bound Bin1 was quantified and shown in (D) (n=3). (E) Effects of CNM-Bin1 on Dyn2 fission activity. 0.5 μM Dyn2 was incubated with SUPER templates in the presence of GTP and indicated Bin1 proteins. The fission activity was determined by sedimentation and release of fluorescent vesicles into the supernatant (n=3). (F) Binding affinity between Bin1 and the PRD of Dyn2. Various concentration of GST-PRD was used to pull down purified Bin1, and then the PRD-bound Bin1 was analyzed. The dissociation constant (Kd) was determined by curve fitting using nonlinear regression and shown with standard error in (G) (n=3). (H) Electron micrographs of Dyn2 together with Bin1 mutants assembled onto liposomes. 1 μM wild-type Dyn2 and 1 μM Bin1 mutants were incubated with 100-nm liposomes for 10 min at room temperature, adsorbed to grids and imaged by negative-stain TEM. Scale, 100 nm. Boxed areas were magnified and shown. The diameter of tubulated membrane was quantified and shown in (I) (n=7). Bar graphs are shown as average ± SD, box plots are shown as median ± min and max value. Data are analyzed with one-way ANOVA. ns, no significance; *P< 0.05; **P< 0.01; ***P< 0.001.

Consistent with the result from *in vitro* membrane tubulation (Fig 2B), expression of GFP-tagged Bin1^Q434X^ and Bin1^K436X^ in C2C12 myoblasts induced tubulation of plasma membrane similar to Bin1^WT^-GFP (Fig S3C). Given that these membrane tubules are highly curved like T-tubules and could be severed by Dyn2 into small vesicles (Chin et al., 2015; Laiman and Liu, 2020), we cotransfected Dyn2-mCherry and Bin1-GFP into C2C12 myoblasts and imaged them under confocal microscopy to examine *in cellulo* Dyn2 activity-regulated by CNM-associated Bin1 mutants (Fig 3A). We then categorized the phenotypes into three classes based on Bin1-induced morphologies; 1) predominately membrane tubules (Tubular); 2) predominately vesicles (Vesicular), 3) relatively equal numbers of tubules and vesicles (Intermediate, Figure 3B). Both Bin1^Q434X^- and Bin1^K436X^-GFP cotransfected cells had significantly fewer cells in the tubular category and more cells in vesicular category (Fig 3B), reflecting diminished inhibition of Dyn2 fission activity by these mutants compared to Bin1^WT^. Live cell imaging enabled us to directly observe the dynamics of Bin1-induced structures in cotransfected cells (Video S1). Bin1^WT^-GFP mainly appeared as plasma membrane derived tubules that are relatively stable overtime (Fig 3C). In contrast, in cells cotransfected with either Bin1^Q434X^-GFP or Bin1^K436X^-GFP, fission events could be seen more frequently with vesicles being cut directly from membrane tubules (arrowheads in Fig 3C).

**Figure 3.**
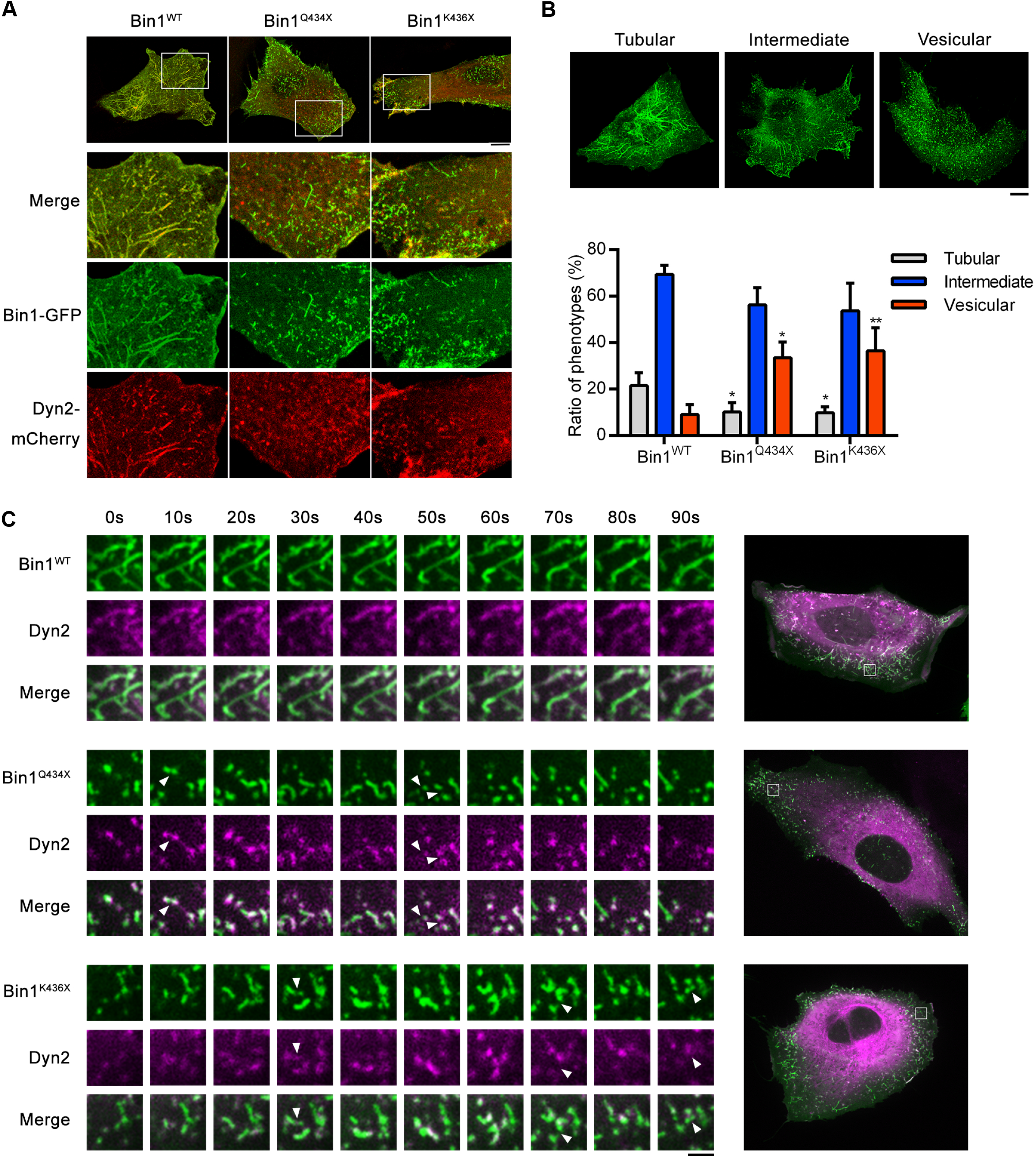
Effects of CNM-Bin1 on Dyn2 fission activity *in cellulo*. (A, B) Morphology of CNM-associated Bin1 mutations co-expressed with Dyn2-mCherry in myoblast. GFP-tagged wild-type or mutant Bin1 was transfected into C2C12 myoblasts together with Dyn2-mCherry. The morphology of Bin1-mediated membrane tubulation was imaged with confocal microscopy. Bottom panels, magnified images from insets in top panels. The Bin1-mediated membrane deformation was categorized into three groups, and the ratio of each population was quantified. Three independent experiment repeats were analyzed with one-way ANOVA and shown in (B). The data was compared with Bin1^WT^ and shown as average ± SD. *P< 0.05, **P< 0.01. (C) Representative Bin1-derived membrane tubules in Dyn2-mCherry co-expressing cells were shown. White arrow heads indicate the fission events occur on Bin1 tubules. Scales, 2 μm.

In summary, CNM-associated Bin1 mutants with SH3 domain truncation retain intact membrane binding and tubulation properties while having decreased interaction with PRD of Dyn2. Consequently, alleviation of Bin1-SH3 inhibition on Dyn2 reversed the overall regulatory effect of Bin1 to promote Dyn2 membrane fission activity instead, demonstrating the pathological mechanism of these CNM-related mutants of Bin1.

### Phosphorylation of Dyn2 PRD attenuates Bin1 binding and inhibition

SH3 domain truncation of Bin1 could relieve Dyn2 inhibition, but it is a pathological condition that appears as a result of CNM-associated Bin1 mutations. We aimed to investigate the mechanism to relieve SH3-PRD inhibition of Dyn2 under physiological condition. Phosphorylation of Dyn1 PRD has been known to regulate its activity and function (Clayton et al., 2010; Reis et al., 2015; Tan et al., 2003), so we hypothesized that Dyn2 might undergo similar regulation. Among the reported phosphorylation sites on the PRD of Dyn2, we chose two sites that are located adjacent to SH3 binding pockets in the C-terminal of Dyn2 PRD, Ser848 and Ser856 (Fig 4A) (Choudhary et al., 2009; Efendiev et al., 2002), then generated phosphomimetic and phosphodeficient mutants of these two sites for the following analysis. Pull-down assays using GST-tagged SH3 domain of Bin1 (GST-SH3) were conducted to determine whether SH3-PRD interactions are altered in these mutants. Indeed, one of the phosphomimetic mutants, Dyn2^S848E^, had lower binding affinity to Bin1-SH3 (Fig 4B and C). This effect was specific to SH3-PRD interaction between Dyn2 and Bin1 as no difference in binding was observed between Dyn2^WT^, Dyn2^S848A^ and Dyn2^S848E^ to the SH3 domain of endophilin (Fig S4A-C).

**Figure 4.**
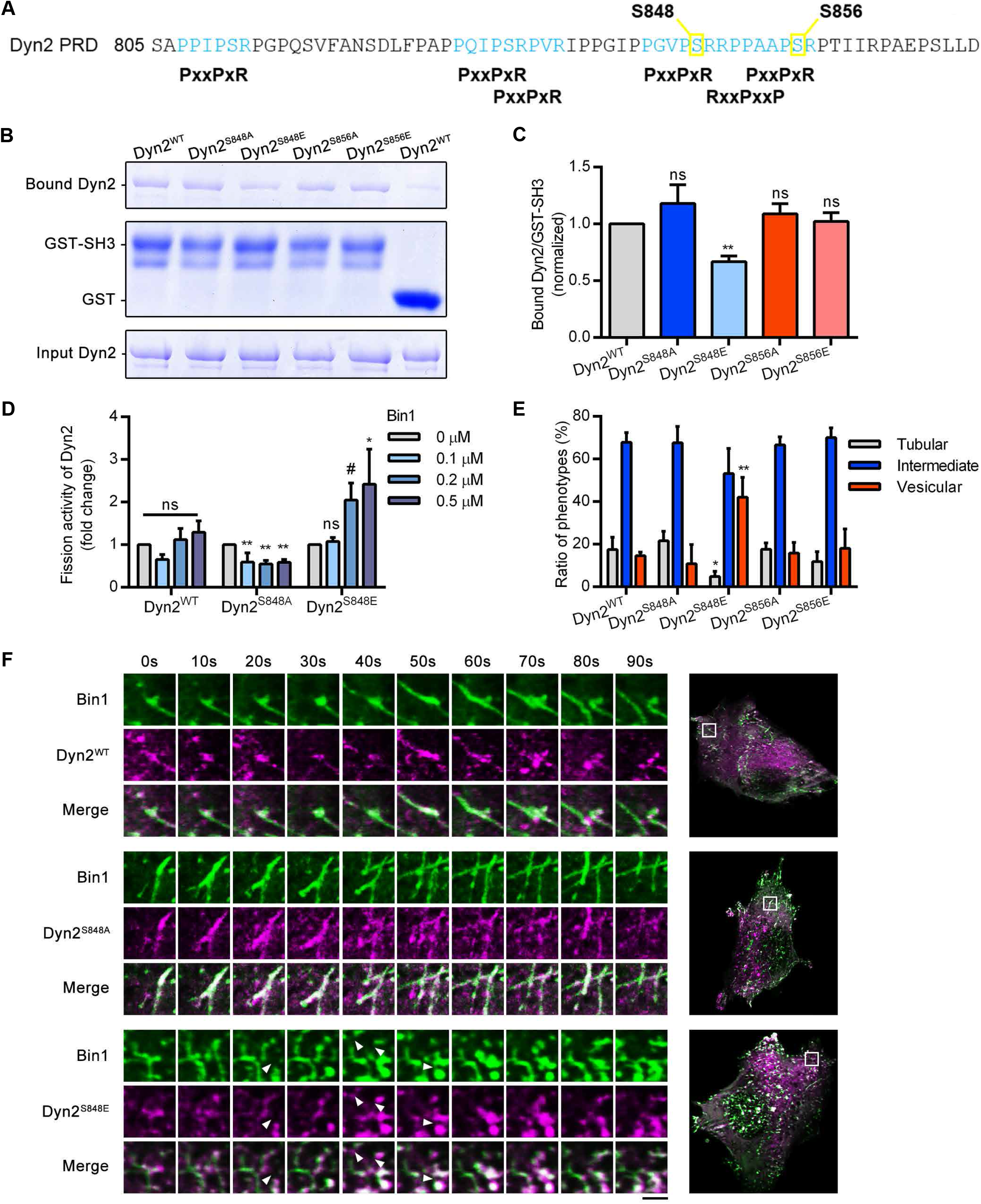
Bin1 enhances the fission activity of phosphorylated Dyn2. (A) The amino acid sequence of Dyn2-PRD. Two potential phosphorylated serine, S848 and S856, were boxed and highlighted in yellow. The SH3-binding pockets, PxxPxR or RxxPxxP, were highlighted in light blue. (B) GST pull-down assay. GST-tagged Bin1 SH3 domain was incubated with indicated Dyn2 mutants. The bound Dyn2 was detected with CBR stain, quantified with imageJ and shown in (C) (n = 3). (D) Fission activity of Dyn2 mutants in the presence of Bin1. 0.5 μM indicated Dyn2 proteins were incubated with SUPER templates in the presence of GTP and indicated Bin1 proteins. The fission activity was determined by sedimentation and release of fluorescent vesicles into the supernatant. The data was shown as fold change relative to the Dyn2 fission activity in the absence of Bin1 (n = 3). (E) Effect of Dyn2 mutants on GFP-Bin1 tubules in myoblasts. Wild-type or indicated mutant Dyn2-mCherry were transfected into myoblasts together with Bin1-GFP. The phenotype of Bin1-GFP tubules was analysed as in Fig 3B. Three independent experiment repeats were analyzed and shown in (E). (F) Time-lapse representative images of Bin1-GFP tubules in cells co-expressing wild-type, S848A or S848E Dyn2-mCherry were magnified and shown. White arrow heads indicate the occurrence of membrane fission. Scale, 2 μm. Data are shown as average ± SD and analyzed with one-way ANOVA. ns, no significance; #P≤ 0.06; *P< 0.05; **P< 0.01.

To determine if weaker SH3-PRD binding would also relieve Dyn2^S848E^ from Bin1-SH3 inhibition, we performed *in vitro* fission assays and confirmed that the fission activity of Dyn2^S848E^ was indeed promoted by Bin1 (Fig 4D). Interestingly, unlike Dyn2^WT^, the non-phosphorylatable mutant Dyn2^S848A^ was inhibited by Bin1, suggesting a shift in the balance between dual stimulatory and inhibitory effects of Bin1. We next compared the sensitivity of these mutants to Bin1-dependent alterations in assembly. Quantification of tubule diameter under TEM showed that membrane tubules derived from Dyn2^S848E^ and Bin1 coassembly on liposome were smaller than those of Dyn2^WT^ and Bin1 in average (Fig S4D and E). Together these data support our hypothesis that phosphorylation of Ser848 in the PRD diminishes Bin1 interactions and thus disrupts its inhibitory effects on Dyn2.

To further validate our hypothesis, we cotransfected mCherry-tagged Dyn2 mutants with Bin1-GFP into C2C12 and classified cells based on Bin1 phenotypes just like in Fig 3B. Compared to Dyn2^WT^, cells cotransfected with Dyn2^S848E^ had significantly more cells predominated by vesicles and fewer cells predominated by tubules (Fig 4E). Cotransfected cells were also observed using live cell imaging to view the dynamics of Bin1 and Dyn2 in cells (Videos S2-4). In Dyn2^WT^ and Bin1 cotransfected cells, Bin1 formed membrane tubules that would elongate and stay as tubules even with Dyn2 colocalization (Video S2 and Fig 4F). Consistent with our *in vitro* results, tubulation was even more pronounced in Dyn2^S848A^ and Bin1 cotransfected cells (Video S3 and Fig 4F). In contrast, Dyn2^S848E^ actively promoted fission of Bin1-tubulated membranes in cotransfected cells (Video S4 and arrowheads in Fig 4F), causing them to be predominated by Bin1 vesicles. Together, these results provide strong evidence that phosphorylation of Dyn2 PRD serves as a mechanism to physiologically attenuate SH3-PRD binding with Bin1 and thus relieve inhibition of Dyn2 fission activity.

### Dyn2-Ser848 phosphorylation modulates clathrin-mediated endocytosis in C2C12 myotubes

After establishing Bin1-SH3 inhibition on Dyn2 and its modulation through PRD phosphorylation, we wondered whether this regulation would affect clathrin-mediated endocytosis (CME). We analyzed endocytic uptake of Alexa488-labelled transferrin (Transferrin-488) in C2C12 myoblasts overexpressing Dyn2^WT^ and various mutants. A Dyn2 mutant with a curvature-generation defect, Dyn2^G537C^, served as a negative control and displayed lower transferrin internalization compared to Dyn2^WT^ (Fig S5A and C). However, no significant difference was detected between either Dyn2^S848A^ or Dyn2^S848E^ transfected myoblasts relative to Dyn2^WT^. Since Bin1 regulation on Dyn2 fission activity happened in a concentration-dependent manner *in vitro* (Fig 1B and S1C), and Bin1 expression was induced after C2C12 myoblasts differentiation (Fig S5E, (Chin et al., 2015)), we proceeded to measure transferrin uptake in C2C12-differentiated myotubes. Under this condition, the phosphodeficient mutant, Dyn2^S848A^, exhibited lower transferrin internalization than Dyn2^S848E^ and Dyn2^WT^ (Fig S5B and D), reflecting stronger restriction by Bin1 on fission activity of Dyn2^S848A^. These results demonstrated the importance of Dyn2-Ser848 phosphorylation on CME in myotubes.

### Dyn2 S848 phosphorylation facilitates GLUT4 internalization in muscle cells

We next investigated the role of binding affinity-regulated Dyn2 fission activity in other dynamin-mediated endocytic pathways, especially in muscle-specific pathways. Glucose transporter type 4 (GLUT4) is predominantly expressed in muscle and adipose tissues where it facilitates glucose uptake (Birnbaum, 1989; Charron et al., 1989; James et al., 1989). In muscle cells, insulin stimulation and exercise trigger GLUT4 translocation from its intracellular storage vesicles to the plasma membrane (Ploug et al., 1998). In the absence of insulin, GLUT4 is reinternalized back into intracellular compartments via Dyn2-mediated endocytosis (González-Jamett et al., 2017; Volchuk et al., 1998). To analyze the effect of Dyn2-Ser848 phosphorylation on endogenous GLUT4 trafficking, we utilized lentiviral overexpression of Dyn2 mutants in C2C12 myotubes followed by subcellular fractionation to separate GLUT4 into two membrane pools by differential centrifugation: plasma membrane-localized GLUT4 pelleted in the heavy membrane fraction (P), whereas intracellular GLUT4 remained in the supernatant (S) (Laiman and Liu, 2020). Myotubes with Dyn2^S848A^ overexpression had a significantly higher ratio of GLUT4 in heavy membrane fraction compared to Dyn2^WT^ and Dyn2^S848E^ infected cells without changing the total amount of GLUT4, indicating less efficient GLUT4 internalization due to Bin1-SH3 inhibition on Dyn2 ^S848A^ (Fig 5A-B, S6A-B).

**Figure 5.**
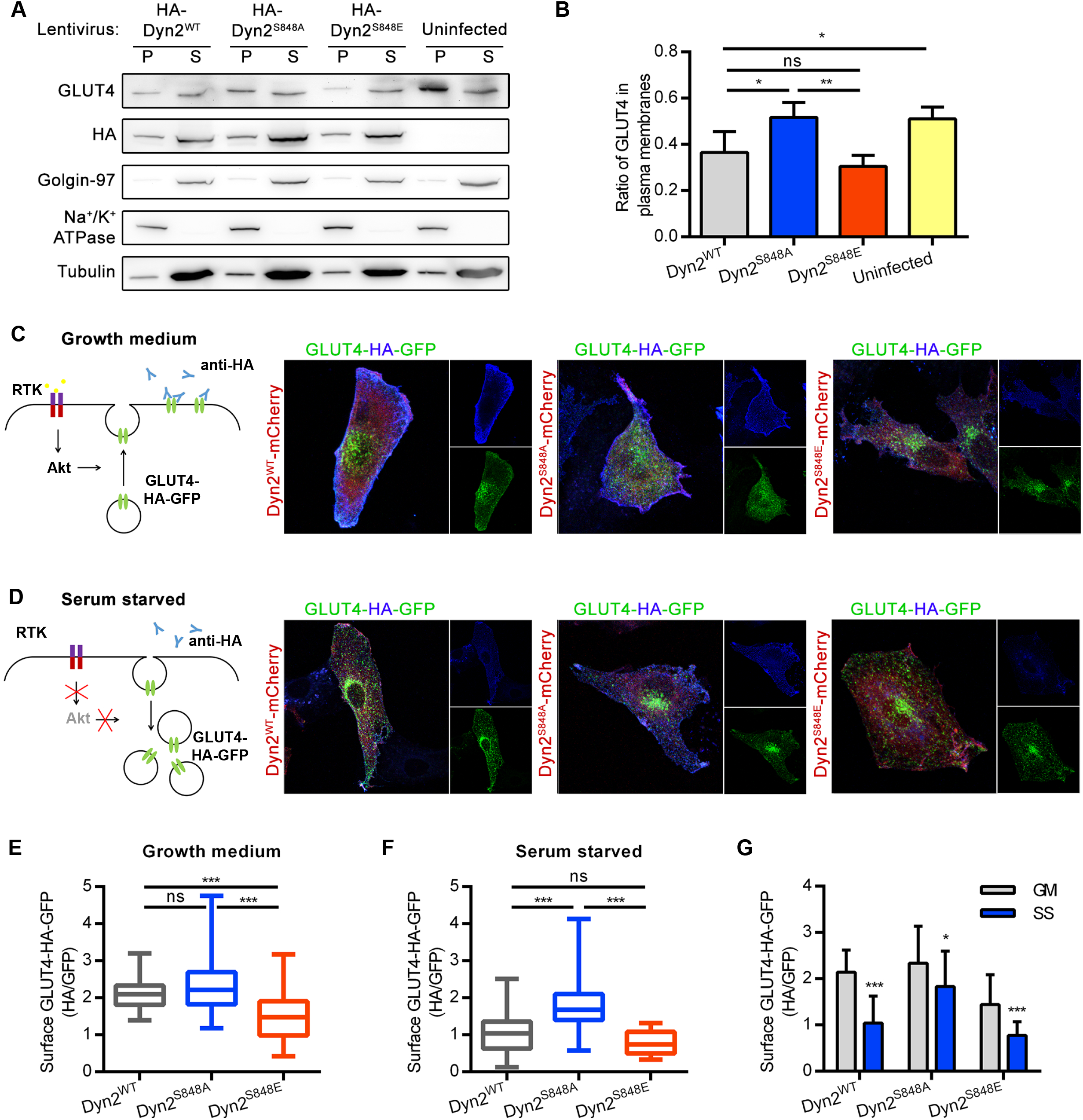
Dyn2^S848E^ promotes the internalization of GLUT4 in myoblasts. (A) Subcellular distribution of endogenous GLUT4 in C2C12 myotubes expressing different Dyn2 mutants. Differentiated C2C12 myotubes were infected with lentiviruses to express indicated HA-Dyn2. After 6 days, these myotubes were subjected to subcellular fractionation to determine the distribution of endogenous GLUT4 by Western blotting. The markers for heavy membrane (Na^+^/K^+^ ATPase, plasma membrane), light membrane (Golgin-97, *trans*-Golgi) and cytosol (tubulin) were used to validate this assay. The ratio of GLUT4 in heavy membrane fraction was quantified and shown in (B) (n = 4). (C, D) Effects of Dyn2-S848 mutations on the insulin-regulated GLUT4-HA-GFP distribution. L6 myoblasts co-transfected with GLUT4-HA-GFP and Dyn2-mCherry mutants were subjected to growth or serum starved media for 2 hours. After immunofluorescent staining with anti-HA antibody without permeabilization and imaging by confocal microscopy, the relative amount of surface GLUT4-HA-GFP was quantified by the fluorescent intensity of HA signaling in blue divided by the total GLUT4-HA-GFP in green. The cartoons illustrate the expected location of GLUT4-HA-GFP under indicated conditions. Three independent experiment repeats were analyzed with one-way ANOVA and shown in (E-G). Bar graphs are shown as average ± SD, box plots are shown as median ± min and max value. ns, no significance; *P< 0.05; **P< 0.01; ***P< 0.001.

To further examine GLUT4 distribution under insulin stimulation, we utilized immunofluorescent staining without permeabilizing the cells to quantify the ratio of surface GLUT4-HA-GFP in L6 myoblasts cultured in growth medium or under serum starvation. When L6 cells were maintained in growth media, GLUT4-HA-GFP could be detected both at cell surface (red signals in Figure S6C) and at intracellular storage vesicles or recycling endosomes (green signals in Fig S6C). GLUT4-HA-GFP distribution was noticeably shifted intracellularly upon incubation in serum-free medium, whereas addition of insulin returned GLUT4 localization to plasma membrane (Fig S6C).

Having established that insulin regulates the localization of GLUT4 in L6 myoblasts, we next analyzed the effects of overexpression of WT and mutant Dyn2 on GLUT4 disposition in growth medium and under serum starvation by quantifying the fluorescent intensity of surface (HA) versus total (GFP) GLUT4-HA-GFP (Fig 5C-G). Under both conditions, significantly higher levels of GLUT4 were located on the cell surface in L6 cells expressing Dyn2^S848A^ compared to Dyn2^S848E^, once again reflecting lower efficiency of GLUT4 internalization. In growth medium, ratio of surface GLUT4 in Dyn2^WT^-cotransfected L6 was similar to that of Dyn2^S848A^ (Fig 5C and E). Interestingly, in growth media, the ratio of surface to total GLUT4 in L6 cells expressing Dyn2^WT^ resemble that of Dyn2^S848A^ expressing cells, whereas after serum starvation, the GLUT4 distribution in Dyn2^WT^ L6 cells switched to resemble that seen in Dyn2^S848E^-expressing cells (Fig 5D and F). These data suggest that in growth media the inhibitory effects of Bin1 on Dyn2 dominate, while after serum starvation Bin1 switches to a stimulatory role. Notably, the expression of either phosphodeficient or phosphomimetic mutants of Dyn2-Ser848 severely reduced the insulin-stimulated GLUT4 translocation in L6 myoblasts (Fig 5G).

To know whether phosphodeficient or phosphomimetic mutants of Dyn2-Ser848 would also affect the recycling pathway, we measured the efficiency of transferrin recycling and found no significant difference in Dyn2-Ser848 mutants (Fig S7). Together, these results establish the importance of Dyn2-Ser848 phosphorylation in modulating the activity of Dyn2 and GLUT4 endocytosis in muscle cells.

### Dyn2-Ser848 phosphorylation is regulated by insulin signaling

While our analysis of phosphorylation mutants of Dyn2 strongly suggest a role for Dyn2-Ser848 phosphorylation in regulating insulin-dependent trafficking in myoblasts, we next sought evidence for phosphorylation of Dyn2 under physiological conditions. For this we performed immunoprecipitation and phosphoserine detection of Dyn2 in L6 myoblasts. Phosphoserine antibody (p-Ser) analysis of Dyn2 phosphorylation status showed that serum starvation slightly elevated the p-Ser signal of HA-Dyn2^WT^ but not HA-Dyn2^S848A^ (Fig 6A and B), indicating increased phosphorylation of Dyn2-Ser848 under this condition.

**Figure 6.**
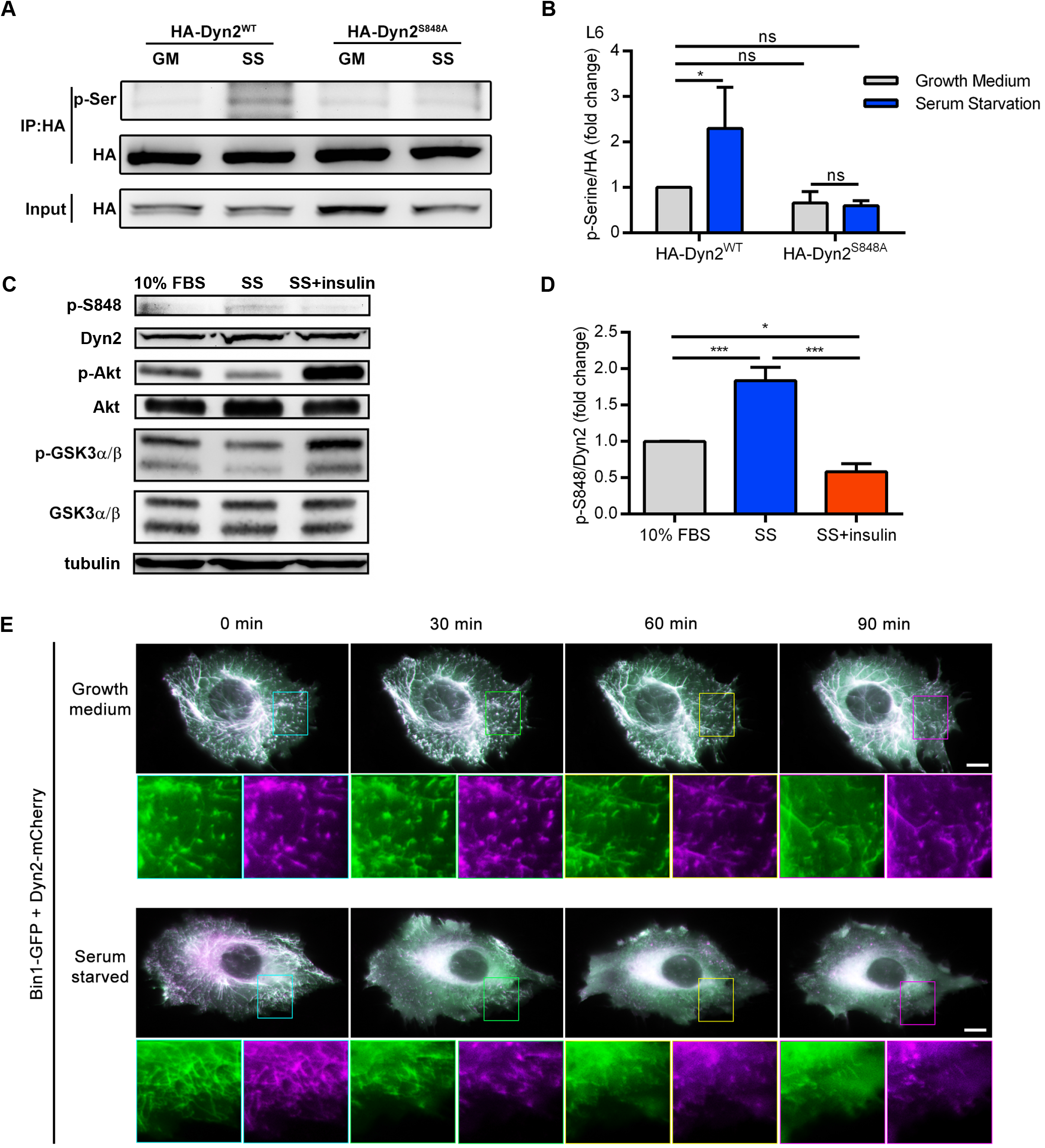
The phosphorylation of Dyn2^S848^ is regulated by insulin signaling. (A) Phosphorylation of Dyn2 in response to serum starvation. HA-tagged Dyn2^WT^ or Dyn2^S848A^ expressed in L6 myoblasts were precipitated by anti-HA antibody with or without serum starvation. Phosphorylated Dyn2 was quantified by anti-phospho-serine antibody and normalized with precipitated HA intensity, compared to HA-Dyn2^WT^ in growth medium and shown in (B). (C) Phosphorylation of endogenous Dyn2 in response to insulin. C2C12 myotubes were subjected to growth medium (10% FBS), 3-hour serum-free medium (SS) or SS followed with 30-min,100 nM insulin stimulation. Cell lysates were then harvested and detected by Western blotting with indicated antibodies. The ratio of phosphorylated Dyn2^S848^ was quantified and shown in (D) (n=3). (E) Time-lapse representative images of Bin1-GFP tubules of cells cultured in growth or serum starved medium. Boxed areas were magnified and shown in the lower panel. Scale, 10 μm. Data are shown as average ± SD and analyzed with one-way ANOVA. ns, no significance; *P< 0.05; ***P< 0.001.

To confirm the phosphorylation of endogenous Dyn2, we generated a phospho-specific antibody recognizing phosphorylated Dyn2-Ser848 (named p-S848) and detected higher p-S848 signal in C2C12 myotube with three hours of serum starvation (Fig S8A). More importantly, the phosphorylation of endogenous Dyn2 was significantly reduced by insulin (Fig 6C and D). Similar to the effect of Dyn2^S848E^ on Bin1 tubule, serum starvation swiftly induced the vesiculation of Bin1 tubules in myoblasts, and this effect could be relieved by one-hour insulin incubation (Fig 6E, S8B and C).

### GSK3α phosphorylates PRD of Dyn2

Utilizing Scansite 4.0 to predict possible kinases of Dyn2S848, we identified glycogen synthase kinase 3α (GSK3α) as a potential candidate. GSK3α is a serine/threonine kinase that is inhibited by insulin-PI3K-Akt signaling through phosphorylation at Ser21, rendering it inactive (Cross et al., 1995; Frame et al., 2001). To test GSK3α as the potential kinase of Dyn2-Ser848, HA-Dyn2^WT^ or HA-Dyn2^S848A^ were coexpressed with either wild type (WT), constitutively active (S21A, GSK3α CA), or kinase inactive (K148A, GSK3α KI) GSK3α in HeLa cells. GSK3α CA significantly enhanced p-Ser signal of HA-Dyn2^WT^, without affecting the extent of phosphorylation of HA-Dyn2^S848A^ (Fig 7A and B). There are two ubiquitously expressed forms of GSK3 in mammals, GSK3α and GSK3β, which share high similarity in their catalytic domains (Woodgett, 1990). We investigated whether GSK3β could also catalyze Dyn2-Ser848 phosphorylation. Unlike GSK3α, co-expression with GSK3β CA produced no particular difference in phosphorylation status between HA-Dyn2^WT^ and HA-Dyn2^S848A^ (Fig 7C, D and S9A), consistent with previous report that Dyn2 was not phosphorylated by GSK3β *in vitro* (Clayton et al., 2010).

**Figure 7.**
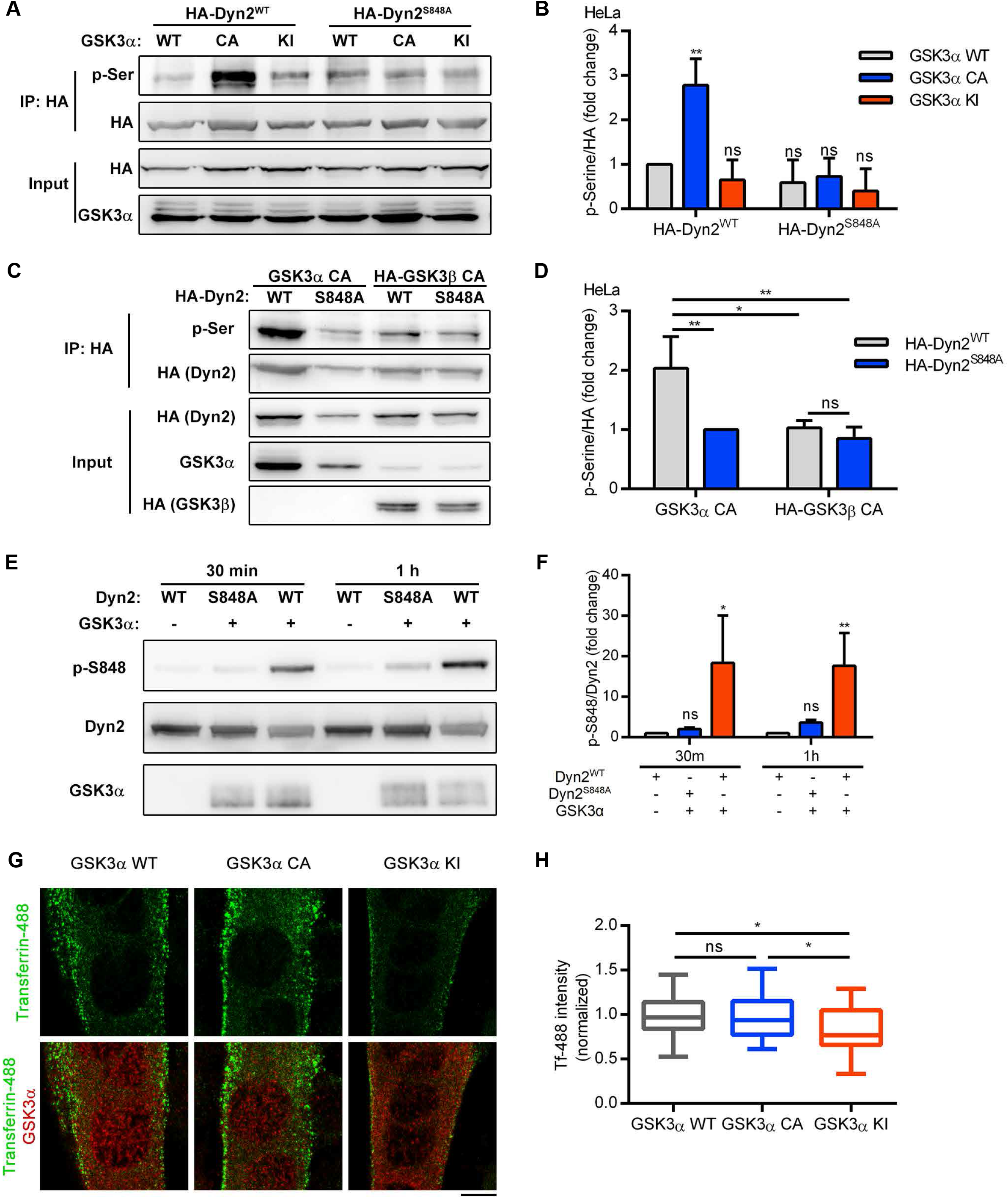
GSK3α phosphorylates the S848 residue of Dyn2. (A) Phosphorylation of Dyn2 in response to GSK3α overexpression. HA-Dyn2^WT^ or HA-Dyn2^S848A^ were co-expressed with WT, CA or KI forms of GSK3α in Hela cells. After precipitation with anti-HA antibody, the phosphorylated Dyn2 was detected with anti-phospho-serine antibody, normalized with HA intensity and shown in (B) (n=3). (C) Isoform specific activity of GSK3 on Dyn2. HA-Dyn2^WT^ or HA-Dyn2^S848A^ were co-expressed with constitutive active (CA) GSK3α or GSK3β in Hela cells. After precipitation with anti-HA antibody, the phosphorylated Dyn2 was detected with anti-phospho-serine antibody, normalized with HA intensity and shown in (D) (n=3). (E) *In vitro* kinase assay. 0.8 μg purified Dyn2^WT^ or Dyn2^S848A^ were incubated with ATP in the presence or absence of 10 ng purified GSK3α. After incubation for 30 min or 1 hour, the phosphorylated Dyn2 was quantified by Western blotting with anti phospho-Dyn2^S848^ antibody. The intensity of p-Dyn2^S848^ was quantified with ImageJ, normalized with signal derived from anti-Dyn2 antibody and shown in (F) (n=3). (G) Effect of GSK3α mutants on transferrin internalization in myotubes. C2C12 myoblasts transfected with indicated GSK3α mutants and differentiated into myotubes were incubated with Alexa-488 labelled transferrin at 37°C for 20 min. Cells were then washed on ice, fixed and imaged under confocal microscopy. Scale, 10 µ m. Fluorescence intensity of internalized transferrin was quantified and shown in (H) (n=25).All data are shown as average ± SD and analyzed with one-way ANOVA. ns, no significance; *P< 0.05; **P< 0.01.

Finally, we performed an *in vitro* kinase assay and demonstrated phosphorylation of Dyn2-Ser848 by recombinant GSK3α (Figures 7E and F), which we confirmed by Mass Spectrometry (Fig S9B). Consistent with the inhibitory effect of Dyn2^S848A^ on transferrin endocytosis, the expression of kinase inactive GSK3α KI also resulted in less transferrin internalization in C2C12 myotubes (Fig 7G and H). Together, these data demonstrate that GSK3α directly phosphorylates Dyn2-Ser848 *in vitro* and is the kinase responsible for phosphorylating Dyn2-Ser848 *in vivo* to promote GLUT4 endocytosis while insulin signaling is switched off.

## Discussion

Activation of the insulin-Akt signaling cascade is known to trigger GLUT4 vesicle exocytosis to promote glucose uptake (Beg et al., 2017; Katome et al., 2003; Leto and Saltiel, 2012). In this study, we provide a scenario for the involvement of the same pathway in modulating GLUT4 endocytosis through GSK3α and Dyn2 in muscle cells (Fig 8).

**Figure 8.**
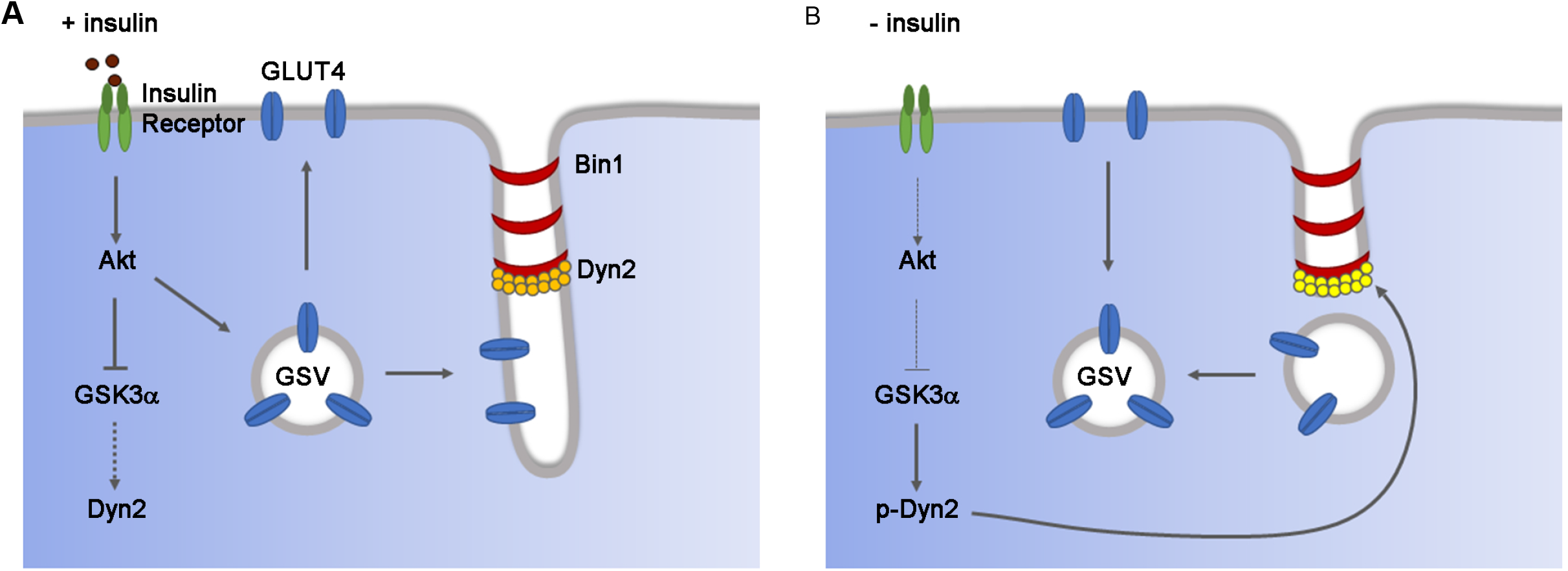
Insulin signaling blocks GLUT4 endocytosis *via* attenuating dynamin-2 fission activity in muscle cells. (A) Insulin signaling promotes the translocation of GLUT4 to muscle surface. Insulin-AKT signaling cascade not only promotes the exocytosis of GLUT4, but also reduces its internalization by suppressing the fission activity of Dyn2. Active Akt phosphorylates thus inactivates GSK3α, resulting in the accumulation of unphosphorylated Dyn2 under inhibitory effect by Bin1. (B) GLUT4 endocytosis is promoted in the absence of insulin signaling. In the absence of insulin signaling, active GSK3α phosphorylates Dyn2^S848^ and relieves the inhibition of Bin1 to accelerate the internalization of GLUT4 from muscle surface.

GLUT4 localizes to sarcolemma and T-tubules in skeletal muscle upon insulin stimulation (Muñoz et al., 1995; Wang et al., 1996). GSK3α is inactivated downstream of Akt after cellular stimulation with insulin or growth factors (Cross et al., 1995). The activity of un-phosphorylated Dyn2 is inhibited by Bin1 interaction thus prevents the internalization of GLUT4 in muscle cells under insulin stimulation (Fig 8A). After insulin is withdrawn and Akt becomes inactive, GSK3α promotes Dyn2-mediated GLUT4 internalization through phosphorylation of Dyn2 PRD, which disturbs interactions with and releases the inhibitory regulation by Bin1 (Fig 7B).

Mammalian GSK3 is composed of two isoforms, GSK3α and GSK3β, that share high similarity but are not functionally redundant (Woodgett, 1990). Knockout of the sole GSK3 homolog in *Drosophila* could be rescued by overexpression of human GSK3β but not human GSK3α (Ruel et al., 1993). Additionally, GSK3β deletion in mice results in embryonic lethality, whereas GSK3α deficient mice are viable and display enhanced insulin sensitivity (Hoeflich et al., 2000; Kerkela et al., 2008; MacAulay et al., 2007). Intriguingly, the PRD of Dyn1 has been shown to be phosphorylated by GSK3β (Clayton et al., 2010), and this modification inhibits Dyn1 activity on endocytosis (Clayton et al., 2009). Reduced Dyn1 inhibition by GSK3β in non-small lung cancer cells give rises to rapid, dysregulated CME, also named adaptive CME (Reis et al., 2015). The isoform-specific regulation between GSK3β and Dyn1 is mirrored in our identification of Dyn2 as specific substrate of GSK3α but not GSK3β. More strikingly, GSK3α phosphorylation positively regulates Dyn2 fission activity while GSK3β phosphorylation leads to Dyn1 inactivation (Fig 8). Due to the diversity of their substrates, inhibition of both GSK3 isoforms might be detrimental as shown by the fatal consequence of GSK3α and GSK3β double deletion in mouse cardiomyocyte (Zhou et al., 2016). The isoform-specific function of GSKα we uncovered demonstrates a potential role for GSK3α as a selective-target for treatment against insulin resistance and type II diabetes.

Dyn2 and Bin1 are interacting partners and their mutations lead to the muscle-specific disease, CNM, despite their ubiquitous expression in human tissues (Bitoun et al., 2005; Bohm et al., 2014; Nicot et al., 2007). The tissue-specific pathogenicity of Bin1 and Dyn2 mutations underlines the importance of their regulations in muscle. Bin1 is highly expressed in skeletal muscle where it serves as a membrane sculptor for the formation of T-tubules (Butler et al., 1997). Earlier studies have shown that CNM-related Bin1 mutations located in its amphiphatic helix, N-BAR, and PI motif reduce membrane tubulation properties of Bin1, which could consequently impair T-tubule biogenesis and maturation (Böhm et al., 2013; Picas et al., 2014; Wu and Baumgart, 2014; Wu et al., 2014). CNM-related Bin1 mutants with SH3 domain truncation display defective binding to Dyn2 but have not been well-characterized for their pathogenic mechanism (Nicot et al., 2007). Here, we illustrate the phenotypes of these Bin1 mutants (Bin1^Q434X^ and Bin1^K436X^) in relation to their effects on Dyn2. We discovered that Bin1 negatively regulates Dyn2 fission activity through SH3-PRD interaction. Decreased Dyn2 binding in Bin1^Q434X^ and Bin1^K436X^ alleviates the inhibitory regulation, tipping the balance toward higher fission activity of Dyn2 and presumably causing excessive T-tubules scission. This is analogous to the gain-of-function in CNM-Dyn2 mutants whose hypermorphic expression triggers T-tubule fragmentation in mouse myotubes and fly muscle (Chin et al., 2015). Due to membrane curvature sensitivity of its PH domain, Dyn2 catalyzes membrane fission only after CCPs have fully matured and when the neck is constricted by other BAR-domain containing proteins in non-muscle cells. By contrast, T-tubules are highly curved (20 - 40 nm in width), thus we speculate that additional mechanisms are needed as a brake to suppress excessive Dyn2 fission. Taken together, disruption in T-tubule homeostasis is the common hallmark of CNM.

The location and function of dynamin is regulated by its binding partners, which may positively or negatively regulate its activity (Hohendahl et al., 2017; Meinecke et al., 2013; Neumann and Schmid, 2013). Dynamin contains several SH3-binding, PxxP motifs in its PRD, enabling it to bind with multiple interacting partners (Solomaha et al., 2005). Here, we investigate the regulation of Dyn2 fission activity by Bin1 and find that weaker binding on Dyn2^S848E^ relieves Bin1 inhibition, indicating that (1) Bin1 regulation of Dyn2 is affinity-dependent, and (2) the SH3 domain of Bin1 likely binds to motifs adjacent to Ser848. A phosphomimetic mutant of this site did not affect binding to endophilin-SH3, indicating endophilin and Bin1 do not bind to the same region of Dyn2. We propose that different SH3-containing proteins exert distinct regulatory effects toward Dyn2 via binding to different PxxP motifs.

A recent study revealed that multimeric interaction is needed for efficient recruitment of Dyn2 to the membrane (Rosendale et al., 2019). Since Bin1 regulates Dyn2 in a concentration dependent manner, and Bin1 binding to Dyn2^S848E^ is weakened but not completely diminished, it is likely that the avidity of oligomerized Bin1 and phosphorylated Dyn2 on membrane is sufficient to recruit Dyn2 to plasma membrane. Due to its high expression level in skeletal muscle, Bin1 could be the main interacting partner to facilitate Dyn2 recruitment in muscle, while simultaneously restraining the fission activity of Dyn2 to preserve T-tubules and regulate GLUT4 trafficking. Interestingly, the phosphorylation of Dyn2^S848^ was reported to inhibit endocytosis in kidney epithelium cells (Efendiev et al., 2002), indicating differential consequences of Dyn2^S848^ phosphorylation among tissues with different SH3 proteins expression. Investigating the outcome of Dyn2 partnership with various SH3-containing proteins both individually and in different combinations would be crucial to uncover the mechanism of Dyn2 regulations in different endocytic pathways and tissues.

The discovery of insulin in 1921 had brought hope and life to millions of people who suffer from diabetes, and the one-hundred-year studies have revealed details of the molecular actions of this amazing hormone in cells and organisms. However, many gaps remain in our understanding of the molecular events downstream of the insulin, both in normal physiology as well as diabetics suffering from insulin resistance. The finding of this work will shed light and bring unique insight into the molecular underpinnings of insulin resistance as well as therapeutic strategy development.

## Materials and Methods

### Protein expression and purification

Human dynamin-2 proteins were expressed in Sf9 insect cells transiently transfected with various constructs and purified via affinity chromatography using the SH3 domain of amphiphysin-2 as reported previously (Liu et al., 2011). Various constructs of human Bin1 isoform 8 were cloned into pGEX-4T-1 or pET30a vectors for expression in E. coli and purified with glutathione Sepharose beads (GE) or Ni-NTA spin columns (Qiagen), respectively, followed by elution as recommended by the manufacturer. Mouse endophilin A1 was purified like Bin1.

### Preparation of lipid templates

Membrane compositions used in this study were DOPC:DOPS:PIP_2_ at 80:15:5 or DOPC:DOPS:PIP2:Rh-PE at 79:15:5:1 for fluorescent-tagged membrane. Liposomes and SUPER templates were prepared as previously described (Neumann et al., 2013). Briefly, for liposome preparation, lipid mixtures were dried, rehydrated in buffer containing 20 mM HEPES (pH 7.5) and 150 mM KCl then subjected to three rapid freeze-thaw cycles followed by extrusion through a 0.1 µ m polycarbonate membrane (Whatman) using Avanti mini extruder. SUPER templates were generated by incubating 2.5 µ m silica beads in a solution containing 100 µ M fluorescent-tagged liposomes and 1 M NaCl for 30 min at room temperature. Excess unbound liposomes were washed four times with water after incubation.

### Fluorescence microscopy

For SUPER templates visualization, images were collected using an inverted Zeiss Axio Observer Z1 microscope equipped with plan-apochromat 100×/1.40 oil DIC M27 objective (Carl Zeiss) and analyzed using ZEN software (Carl Zeiss). For cells fixed with 4% formaldehyde and permeabilized with 0.1% saponin, images were acquired using a confocal microscope (LSM700, Carl Zeiss) equipped with plan-apochromat 63×/1.40 oil DIC M27 objective (Carl Zeiss) and processed using ZEN software (Carl Zeiss).

### Fission and SUPER template tubulation assays

Fission activity of Dyn2 was measured as previously described (Chin et al., 2015). Briefly, SUPER templates were incubated with 0.5 µ M Dyn2, 1 mM GTP and indicated concentrations of Bin1 variants in assay buffer (20 mM HEPES (pH 7.5), 150 mM KCl, and 1 mM MgCl_2_) for 30 min at room temperature. The mixtures were spun down at low speed (260g) in swinging bucket rotor to separate SUPER template from the released vesicles. Fluorescence intensity was measured in a plate reader (RhPE excitation = 530/25 nm bandwidth and emission = 580/25 nm bandwidth). The fission activity was expressed as the fraction of released lipids from total SUPER templates. For visualization of membrane tubulation, SUPER template were slowly added in an evenly distributed manner to solution containing 20 mM HEPES (pH 7.5), 150 mM KCl and 0.5 µ M wild-type or mutant Bin1 on BSA coated eight-chamber slides. After 10 min incubation at room temperature, images of SUPER templates were acquired using Zeiss inverted fluorescent microscope Axio Observer Z1 with a 100×/1.40-NA oil-immersion objective with CCD monochrome Photometrics CoolSNAP HQ.

### Liposome binding, transmission electron microscopy (TEM) and GTPase activity assays

To analyze membrane binding ability, 0.5 µ M Dyn2 and/or Bin1 were incubated with 150 µ M, 100 nm liposomes at 37°C for 10 min. Pellets containing liposome-bound proteins were separated from soluble fractions containing unbound proteins (supernatant) by centrifugation at 15,000g and resuspended in assay buffer (20 mM HEPES (pH 7.5), 150 mM KCl). Pellet and supernatant were run on SDS-PAGE, stained with Coomassie Blue and protein intensities were quantified using ImageJ. Membrane binding ability was expressed as the fraction of membrane-bound proteins divided by total proteins.

To visualize protein assembly on membrane, 1 µ M Dyn2 and/or 1 µ M Bin1 were incubated with 25 µ M liposome for 10 min at 37°C. The mixture was then adsorbed onto the surface of glow-discharged (30 s), carbon film-supported grids (200 mesh copper) for 5 min followed by 2 min negative-staining with 2% uranyl acetate. Images were captured with a Hitachi H-7650 electron microscope operated at 75 kV and a nominal magnification of 150,000x. Liposome tubules diameter were measured using ImageJ.

For GTPase activity assay, 0.5 µ M purified Dyn2 was incubated with 0.5 µ M Bin1 or Endophilin in reaction mixture containing 150 µ M liposomes, 20 mM HEPES (pH 7.4), 150 mM KCl, 1 µ M MgCl_2_ and 1mM GTP at 37°C. Liposome-stimulated GTPase activity of Dyn2 was measured as a function of time using a colorimetric malachite green assay which monitors the release of inorganic phosphate (Leonard et al., 2005).

### GST pull down assay

To analyze SH3-PRD binding affinity, purified His-Bin1 proteins were first incubated with 100 nm liposomes containing 5% PI(4,5)P_2_ to alleviate the autoinhibition between SH3 domain and PI motif before being incubated on 1:2 ratio with GST or GST-tagged PRD of Dyn2 immobilized on glutathione beads for 30 min at room temperature. After washing, the beads were boiled in sample buffer and bound protein were detected by SDS-PAGE followed by Coomassie Blue staining. ImageJ software was used to quantify protein bands intensity of bound Bin1 relative to GST-PRD. Pull down assay for purified Dyn2 mutants with GST-tagged SH3 domain of Bin1 was performed in a similar manner without the presence of 100 nm liposomes.

### Cell culture, transfection and lentiviral infection

Mouse-derived C2C12 myoblasts (ATCC, CRL-1772) were cultured in growth medium (GM) composed of high glucose (4.5 mg/mL) DMEM supplemented with 2mM L-glutamine, 1 mM sodium pyruvate, antibiotics, and 10% fetal bovine serum (Gibco). Upon reaching 90% confluency, C2C12 differentiation was induced by replacing GM with differentiation medium (DM) composed of high glucose (4.5 mg/mL) DMEM supplemented with 1 mM sodium pyruvate, antibiotics, and 2% horse serum (Gibco). The first time cells were incubated in DM was regarded as day 0 of differentiation. Rat-derived L6 myoblasts (ATCC, CRL-1458) were cultured in low glucose (1.0 mg/mL) DMEM supplemented with antibiotics and 10% fetal bovine serum (Gibco). For transfection, cells at 70% confluency were transfected with plasmid of interest using Lipofectamine 2000 (Invitrogen) or *Trans*IT-X2 (Mirus Bio) as suggested by the manufacturers. For lentiviral infection, C2C12 myoblasts at 50% confluency were infected with viruses in the presence of 12 µ g/ml polybrene followed by puromycin (2µg/ml) selection 24h post-infection for 3 days before DM replacement. 10 µ g/ml doxycycline were added to day 3 differentiated C2C12 and GLUT4 distribution in cells was analyzed at day 6 of differentiation. List of plasmids used for transfection and lentiviral generation in this study could be found in Table S1.

### Live cell imaging

For time-lapse microscopy, cells were imaged with spinning disc confocal microscope or lattice-light sheet microscope (Chen et al., 2014). For spinning disc confocal microscope, C2C12 myoblasts were seeded on glass-bottom dishes and cotransfected with Bin1-GFP and Dyn2-mCherry. Cells were placed at 37°C in imaging medium (phenol-red free DM with 20 mM HEPES, pH 7.4, 50 μg/ml ascorbic acid, and 10% FBS) and images were captured with 200 ms (488 nm laser) and 400 ms (561 nm laser) exposure and 5 s interval time using Carl Zeiss Cell Observer SD equipped with plan-apochromat 100 ×/1.40 oil DIC M27 objective (Carl Zeiss). For lattice light-sheet microscopy, C2C12 myoblasts cotransfected with Bin1-GFP and Dyn2-mCherry were immersed in an imaging medium filled chamber. Images were illuminated by exciting each plane with a 488 nm laser at 12.56 nano-Watt (nW) (at the back aperture of the excitation objective) and 561 nm laser at 37.2 nW, with an excitation outer/inner numerical aperture parameters of 0.55/0.44. Orthogonal to the illumination plane, the water immersed objective lens (Nikon, CFI Apo LWD 25XW, 1.1 NA, 2 mm WD) mounted on a piezo scanner (Physik Instrumente, P-726 PIFOC) is used to collect the fluorescence signal, which is then imaged through an emission filter (Semrock Filter: FF01-523/610-25) onto sCMOS camera (Hamamatsu, Orca Flash 4.0 v2 sCOMS) through a 500 mm tube lens (Edmund 49-290, 500 mm FL/50 mm dia; Tube Lens/TL) to provide the 63X magnification observation. And then the cells were imaged through entire cell volumes at 5 s intervals by using the sample piezo (Physik Instrumente, P-622 1CD) scanning mode. Finally, raw data were then deskewed and deconvoluted using GPU_decon_bin (Chen et al., 2014). And using Amira software (Thermo Fisher) to display the 3D images.

### Transferrin uptake assay

Transferrin uptake assay was performed as previously reported (Liu et al., 2008). Briefly, C2C12 transfected with mCherry-tagged Dyn2 mutants and seeded on coverslips were incubated with 5 μg/ml Alexa-488-conjugated transferrin for 10 min (myoblasts) or 20 min (myotubes) at 37°C. For recycling efficiency, 30 min incubation with transferrin-488 was followed by 30 min incubation in growth medium without transferrin-488. Cells were washed with ice-cold PBS then with acid buffer (150mMNaCl and 150 mM glycine, pH 2.0) repeatedly. Afterward cells were fixed with 4% FA and mounted on glass slides using Fluoromount-G mounting medium (Southern Biotech). Images of the cells were acquired using confocal microscopy and analyzed with ZEN software (Carl Zeiss).

### Subcellular fractionation and analysis of surface:total GLUT4 ratio

Subcellular fractionation to analyze GLUT4 distribution was done as previously described (Laiman and Liu, 2020). Briefly, lentiviral infected C2C12 myotubes were homogenized in ice-cold HES-PI buffer (255 mM sucrose, 20 mM HEPES (pH 7.4), 1 mM EDTA, 2x cOmplete protease inhibitors (Roche)). The lysates were then cleared by centrifugation at 1,000 g for 5 min at 4°C. After centrifugation at 16,000 g for 20 min at 4°C, the supernatant was separated from pellet and subjected to TCA precipitation for 1 h at 4°C. Both pellet and supernatant were loaded to SDS-PAGE followed by immunoblotting to detect GLUT4 and markers for heavy membrane, light membrane and cytosol. All antibodies used in this study are listed in Table S2.

Insulin-stimulated HA-GLUT4-GFP localization was done according to previous report (Zeigerer, Lampson et al., 2002). Briefly, L6 were seeded on fibronectin coated coverslips then cotransfected with Dyn2 mCherry and GLUT4-HA-GFP. Cells were incubated in growth medium or serum starved for 2 h prior to fixation and subjected to immunofluorescent staining with anti HA without permeabilization. Cell was imaged under LSM700 confocal microscope (Carl Zeiss) equipped with plan-apochromat 63×/1.40 oil DIC M27 objective (Carl Zeiss) and analyzed using ZEN software (Carl Zeiss). The relative amount of surface GLUT4-HA-GFP was quantified by the fluorescent intensity of HA signal divided by the total GFP signal.

### Immunoprecipitation

For immunoprecipitation, cells were washed with ice-cold 1x PBS added with 1.5 mM Na3VO4 and 50mM NaF before being lysed in lysis buffer (1X PBS, 1% Tx-100, 1.5 mM Na3VO4, 50mM NaF, PhosSTOP (Roche) and cOmplete protease inhibitors (Roche)). The lysates were centrifuged at 16,000g for 10 min at 4°C. The supernatant was then incubated with anti-HA Agarose (Sigma-Aldrich, Cat# A2095) for 1 h at 4°C on a rotator. The beads were washed before being boiled in sample buffer for the following SDS-PAGE and immunoblotting. All antibodies used in this study are listed in Table S2. Band intensities were quantified using ImageJ.

### *In vitro* kinase assay

For in vitro phosphorylation of Dyn2, recombinant Dyn2 was purified from Sf9 as mentioned above, active recombinant GSK3α was purchased (Millipore, Cat# 14-492). 0.8 µg Dyn2WT or Dyn2S848A was incubated with 10 ng GSK3α in kinase buffer (50 mM HEPES (pH 7.4), 15 mM MgCl2, 200 µ M Na3VO4, 100 µ M ATP). The total reaction volume was 30 µ l. Following 30 min or 1 h incubation at 30°C, reaction was stopped by adding sample buffer and was boiled at 90°C before subjected to SDS-PAGE and Western blot analysis with phospho-S848-specific (p-S848) antibody. p-S848 antibody is homemade polyclonal antibody through immunization against phosphoSer848 on Dyn2 using synthetic phosphopeptide with following sequence: 844 PGVP(pSer)RRPPAAPSRC 858, and purified with affinity column.

### Mass spectrometry

For analysis of Dyn2 phosphorylation status, recombinant Dyn2 phosphorylated by GSK3α in vitro as mentioned above was subjected to SDS-PAGE. The gel was then stained with Coomassie Blue and bands containing Dyn2 were cut from the gels as closely as possible. The gel pieces were destained, followed by reduction, alkylation and dehydration before undergoing in-gel digestion with trypsin. The resulting peptides were extracted and went through desalting in C18 column before LC-MS/MS analysis. Peptides were separated on an UltiMate 3000 LCnano system (Thermo Fisher Scientific). Peptide mixtures were loaded onto a 75 μm ID, 25 cm length C18 Acclaim PepMap NanoLC column (Thermo Scientific) packed with 2 μm particles with a pore of 100 Å . Mobile phase A was 0.1% formic acid in water, and mobile phase B was composed of 100% acetonitrile with 0.1% formic acid. A segmented gradient in 50 min from 2% to 40% solvent B at a flow rate of 300 nl/min. LC-MS/MS analysis was performed on an Orbitrap Fusion Lumos Tribrid quadrupole mass spectrometer (Thermo Fisher Scientific). Targeted mass spectrometry analysis was performed. Mass accuracy of <5 ppm, and a resolution of 120,000 at m/z=200, AGC target 5e5, maximum injection time of 50 msec) followed by HCD-MS/MS of the focused on m/z 633.8338 (2+) and m/z 422.8916 (3+). High-energy collision activated dissociation (HCD)-MS/MS (resolution of 15,000) was used to fragment multiply charged ions within a 1.4 Da isolation window at a normalized collision energy of 32. AGC target 5e4 was set for MS/MS analysis with previously selected ions dynamically excluded for 180 s. The MS/MS spectra of pS848 phosphopeptide were manually identified and checked.

### Statistical analysis

Quantitative data in this study are expressed as mean ± SD from at least three independent experiments. GraphPad Prism 8.0 was used for statistical analysis and graphs generation. All data were analyzed with one-way ANOVA followed by Dunnett’s or Tukey’s *post hoc* test, or Student’s t test. Statistical significance was defined using GraphPad Prism 8.0. P < 0.05 was considered statistically significant, indicated as *P< 0.05; **P< 0.01; ***P< 0.001.

## Acknowledgement

We thank the mass spectrometry technical research services from NTU Consortia of Key Technologies and NTU Instrumentation Center, as well as the electron microscopy (EM) core in NTU for their technical support. We are grateful to Dr. Timothy E. McGraw for sharing HA-GLUT4-GFP plasmid and Dr. Sandra Schmid for critical reading and helpful advice on this paper. This work was supported by Ministry of Science and Technology grants MOST 109-2628-B-002-051 and NTUH translational medicine grant to Y.-W. Liu.

## Author contribution

All authors participated in the experimental design; J. Laiman, J. Loh and YL performed major experiments and analyzed data; WT and BC assisted in live cell imaging and data interpretation; J. Laiman and YL wrote the manuscript; BC, YC, LC and YL supervised the project.

